# Expansion of highly interferon-responsive T cells in early-onset Alzheimer’s disease

**DOI:** 10.1101/2023.09.26.559634

**Authors:** Daniel W. Sirkis, Caroline Warly Solsberg, Taylor P. Johnson, Luke W. Bonham, Alexis P. Oddi, Ethan G. Geier, Bruce L. Miller, Gil D. Rabinovici, Jennifer S. Yokoyama

**Affiliations:** Memory and Aging Center, Department of Neurology, Weill Institute for Neurosciences, University of California, San Francisco, San Francisco, CA 94158, USA; Pharmaceutical Sciences and Pharmacogenomics Graduate Program, University of California, San Francisco, San Francisco, CA 94158, USA; Center for Alzheimer’s and Related Dementias, National Institutes of Health, Bethesda, MD 20892, USA; DataTecnica LLC, Washington, DC 20037, USA; Department of Radiology and Biomedical Imaging, University of California, San Francisco, San Francisco, CA 94158, USA; Transposon Therapeutics, Inc., San Diego, CA 92122, USA; Global Brain Health Institute, University of California, San Francisco, San Francisco, CA 94158, USA and Trinity College Dublin, Dublin, Ireland

**Keywords:** Alzheimer’s disease, Early-onset Alzheimer’s disease, Tauopathy, Peripheral blood mononuclear cells, Cerebrospinal fluid, T cells, CD4 T cells, Interferon, Interferon signaling-associated gene, Single-cell RNA-sequencing, Droplet digital PCR

## Abstract

**INTRODUCTION:** Altered immune signatures are emerging as a central theme in neurodegenerative disease, yet little is known about immune responses in early-onset Alzheimer’s disease (EOAD).

**METHODS:** We examined single-cell RNA-sequencing (scRNA-seq) data from peripheral blood mononuclear cells (PBMCs) and droplet digital (dd)PCR data from CD4 T cells from participants with EOAD and clinically normal controls.

**RESULTS:** We analyzed ~182,000 PBMCs by scRNA-seq and discovered increased interferon signaling-associated gene (ISAG) expression and striking expansion of antiviral-like ISAG^hi^ T cells in EOAD. We isolated CD4 T cells from additional EOAD cases and confirmed increased expression of ISAG^hi^ marker genes. Publicly available cerebrospinal fluid leukocyte scRNA-seq data from late-onset mild cognitive impairment and AD also revealed increased expression of interferon-response genes.

**DISCUSSION:** ISAG^hi^ T cells, apparently primed for antiviral activity, are expanded in EOAD. Additional research into these cells and the role of heightened peripheral IFN signaling in neurodegeneration is warranted.

## 1 BACKGROUND

Approximately 5–10% of the ~7 million Americans living with Alzheimer’s disease (AD) [1] experience symptom onset prior to age 65 [2]. In this early-onset form of AD (EOAD), affected individuals are more likely to experience an aggressive clinical course, have an atypical clinical syndrome, encounter delays in diagnosis, and experience unique social disruptions due to their relatively young age [2]. The vast majority (≥90%) of EOAD cases are not inherited in an autosomal-dominant manner, and for these individuals, we understand relatively little about the genetic and other biological factors underpinning disease risk.

Recent reports using single-cell RNA-sequencing (scRNA-seq) have highlighted changes in peripheral blood and cerebrospinal fluid (CSF) leukocyte populations in AD [3], Lewy body dementia [4], familial tauopathy [5], and during aging [6]. To our knowledge, however, a global, unbiased scRNA-seq analysis of peripheral blood mononuclear cells (PBMCs) in EOAD has not been reported. Using scRNA-seq, we now find evidence for marked expansion of a small population of recently characterized CD4 T cells expressing very high levels of interferon (IFN) signaling-associated genes (ISAG^hi^ T cells) in EOAD. Remarkably, a CD4 T-cell subtype that appears to be highly similar to ISAG^hi^ T cells–with a similar antiviral gene expression signature– is potently expanded in the CSF in the context of viral encephalitis [7], suggesting that EOAD-expanded ISAG^hi^ T cells have antiviral properties. Adding to the weight of evidence for augmented peripheral IFN signaling in EOAD, we also observe expansion of a rare natural killer (NK) cell population previously associated with heightened IFN signaling [8].

Beyond changes in cell-type abundance, we report global upregulation of IFN signaling genes across additional lymphoid and myeloid PBMC types in EOAD. In addition, by analyzing a publicly available scRNA-seq dataset of CSF leukocytes derived primarily from individuals with mild cognitive impairment (MCI) and late-onset AD (LOAD) [3], we find striking upregulation of the same IFN signaling pathways in CD4 T cells in late-onset disease. These findings suggest at least partially conserved IFN responses between EOAD and LOAD. Finally, we find consistent upregulation in the hippocampus of a suite of IFN response genes in a mouse model of familial EOAD. Collectively, our findings indicate that dysregulation of IFN-related genes extends from the peripheral blood and CSF to the brain in AD and suggest a novel role for a population of unusual, antiviral-like T cells in EOAD.

## 2 METHODS

### 2.1 Overview

After obtaining informed consent, PBMCs from study participants (Table 1) at the University of California, San Francisco Memory and Aging Center (MAC) were analyzed by scRNA-seq essentially as described [5]. Raw sequencing reads were aligned to GRCh38-2020-A and feature-barcode matrices generated using Cell Ranger (v7.1.0) with intronic reads excluded. Cluster proportions were determined for individual samples by dividing the number of cells in a given cluster by the total number of cells in clusters representing all PBMCs, all T cells, or all CD4 T cells (after quality control) for each individual. Statistical differences in cell-type abundances by diagnosis were assessed via linear modeling, controlling for age and sex. Additional details, including bioinformatic and experimental methods, are described in the Supplementary Methods document.

**Table 1.**
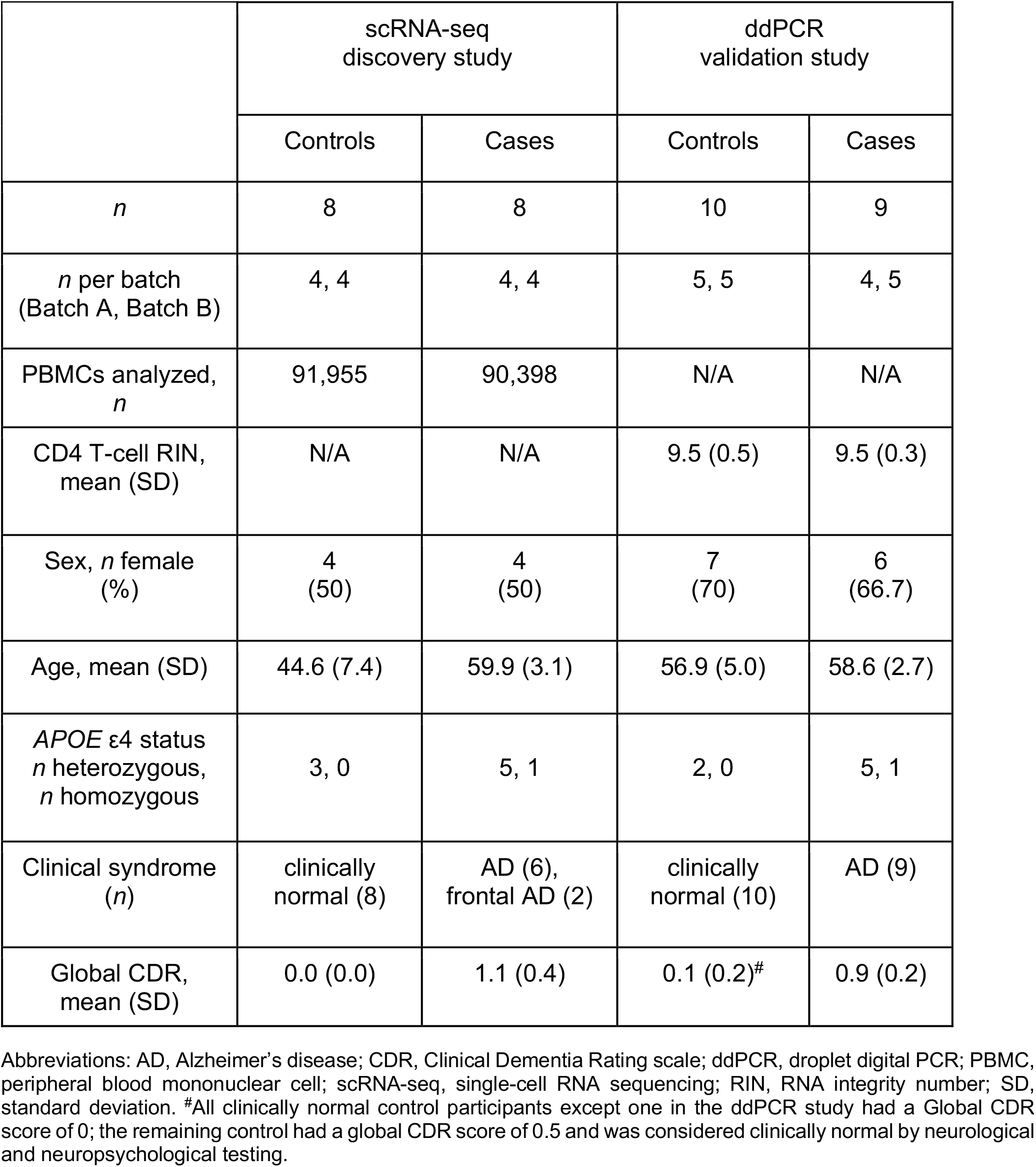
Demographic and experimental information for samples used in scRNA-seq and ddPCR studies.

### 2.2 Clinical assessment

Study participants underwent a multistep screening prior to an in-person clinical assessment at the MAC that included a neurologic exam, detailed cognitive assessment, medical history, and family history for neurodegenerative disease [9]. Study partners were interviewed about the participant’s functional abilities. A multidisciplinary team consisting of a neurologist, a neuropsychologist, and a nurse reviewed all participant clinical information and established diagnoses for cases according to consensus criteria for AD [10–12] or frontal AD [13]. All EOAD cases were diagnosed with probable AD and had at least one positive biomarker consistent with AD. In particular, 10/13 cases had evidence of amyloid and tau positivity (obtained via PET imaging or assessment of CSF amyloid-β_42_ and phospho-tau181 levels) in addition to neurodegeneration (via structural MRI; A+T+N+), while 3/13 cases had evidence of neurodegeneration (N+). The mean (SD) age of first abnormal diagnosis for all participants with EOAD was 58.5 (3.0) and the range was 54–62. All control participants had a normal neurologic exam, and all controls except one in the ddPCR study had a global Clinical Dementia Rating (CDR) [14] scale score of 0; the remaining control participant, who was diagnosed as clinically normal, had a CDR score of 0.5 due to subjective memory complaint. This participant also reported depressive symptoms. Sensitivity analysis of the ddPCR data after exclusion of this individual indicated that the results remained similar or unchanged. All participants screened negative for disease-causing pathogenic variants in established genes for AD and frontotemporal lobar degeneration, which also causes early-onset dementia.

## 3 RESULTS

### 3.1 Identification of an expanded T-cell subtype in EOAD

After QC filtering, clustering of ~182,000 PBMCs generated 19 primary clusters consisting of all expected PBMC types. Comparison of relative cluster abundance in EOAD cases vs. controls revealed a single cluster (cluster 15) that was robustly expanded in EOAD (Figure 1A). Expression of marker genes indicates that cluster 15 is a subtype of CD4 T cell (supplementary Figure S1A). Quantification of cluster 15 abundance relative to either all PBMCs, all T cells, or all CD4 T cells revealed significant expansion in EOAD that was driven primarily by females (Figure 1B and C). To determine what type of CD4 T cell cluster 15 represents, we subsetted all T cells and reclustered them separately from all other cell types. Reclustering revealed this cell type in sub-cluster 11, which expresses uniquely high levels of IFN-signaling genes *MX1* and *IFI6* relative to all other T cells (Figure 1D). As expected, sub-cluster 11 was also significantly expanded in EOAD relative to controls (Figure 1D).

**Figure 1.**
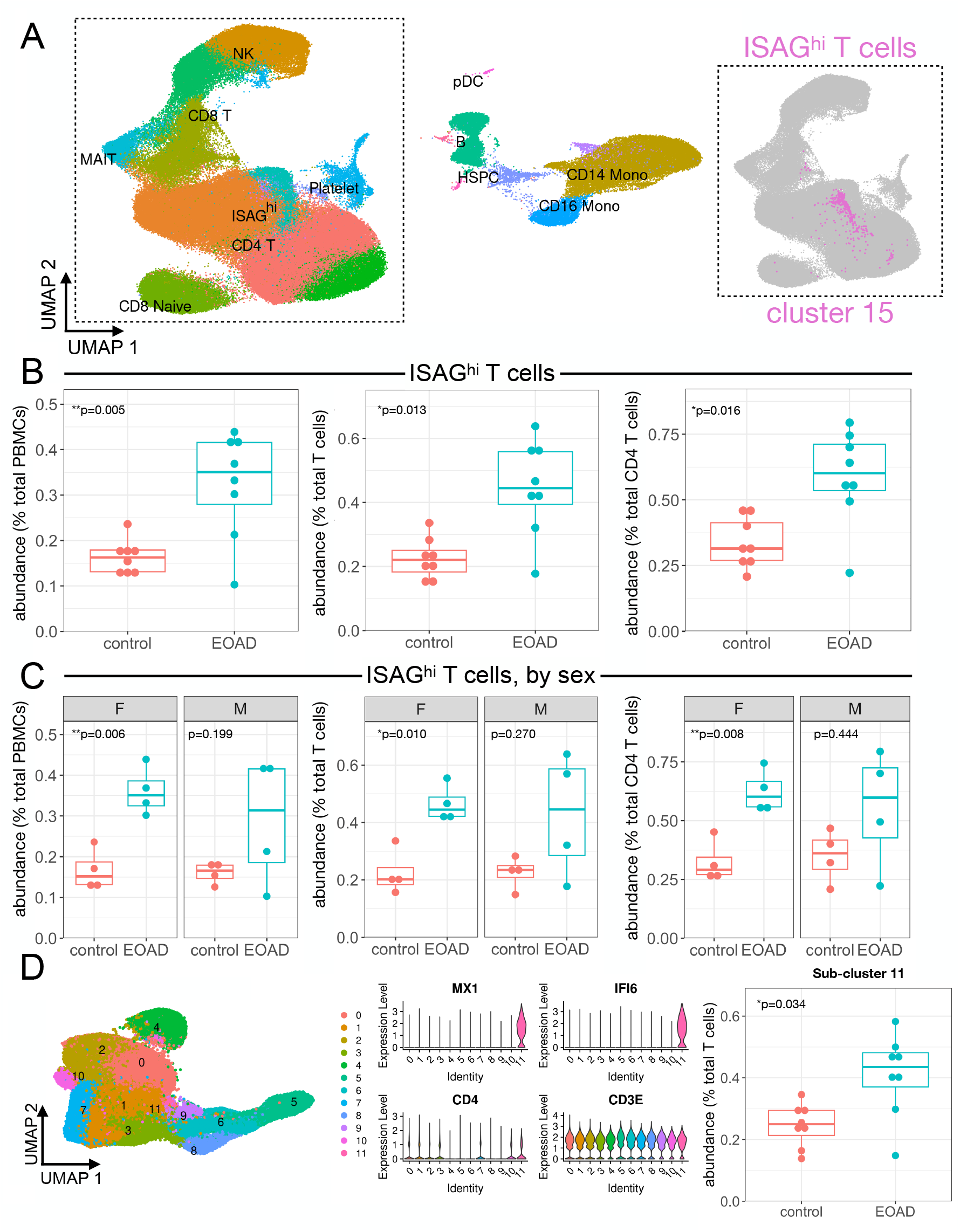
Expansion of ISAG^hi^ T cells in EOAD characterized by scRNA-seq. **A**, Uniform manifold approximation and projection (UMAP) plot of ~182,000 PBMCs from EOAD cases and cognitively normal controls, colored by cluster identity. Major cell types are labeled within the plot. The inset (right) shows the primary T-cell grouping displayed in gray, with the ISAG^hi^ T-cell cluster shown in magenta. **B**, ISAG^hi^ T-cell abundance is quantified relative to all PBMCs (left; *P* = 0.005), all T cells (middle, *P* = 0.013), and all CD4 T cells (right; *P* = 0.016). **C**, Stratifying by sex, ISAG^hi^ T-cell relative abundance is significantly increased in EOAD only in females, expressed as a percentage of PBMCs (left, *P* = 0.006), T cells (middle, *P* = 0.01), and CD4 T cells (right, *P* = 0.008). **D**, Reclustering of all T cells (left) generates a T-cell subcluster (11) representing ISAG^hi^ T cells, which express high levels of marker genes *MX1* and *IFI6*, in addition to T-cell markers *CD4* and *CD3E* (middle). Quantification of the ISAG^hi^ subcluster relative to all T cells again indicates a significant increase in EOAD cases (right, *P* = 0.034).

### 3.2 Characterization of the expanded cell type as ISAG^**hi**^ **T cells**

What is the precise identity of this subset of CD4 T cells? Recent literature using scRNA-seq to analyze human leukocyte populations has revealed two poorly understood cell types: ISAG^hi^ T cells, detected in peripheral blood [15], and antiviral CD4 T cells, detected in CSF [7]. Antiviral CD4 T cells were so named due to their marker gene expression and robust expansion in the CSF in the context of viral encephalitis [7]. Comparison of all marker genes for our sub-cluster 11 to the top 200 marker genes for CSF antiviral CD4 T cells revealed highly statistically significant overlap (*P* = 6.5 × 10^−14^; supplementary Table S1) [16]. Moreover, all of the 12 most-significant marker genes originally reported for ISAG^hi^ T cells [15] are also top marker genes of our sub-cluster 11 and of antiviral CD4 T cells. Therefore, from here on we refer to the EOAD-expanded CD4 T cells as ISAG^hi^ T cells.

### 3.3 Analysis of ISAG^hi^ T-cell abundance in additional samples and datasets

ISAG^hi^ T-cell abundance was consistent across scRNA-seq batches (supplementary Figure S1B) and was not driven by *APOE* ε4 status (supplementary Figure S1C). Moreover, although our control samples came from participants with a younger mean age (Table 1), there was no relationship between age and ISAG^hi^ T-cell abundance (supplementary Figure S1D). To increase the sample size of our scRNA-seq dataset, we included seven additional control PBMC samples previously characterized by scRNA-seq [5]. We found that the expansion of ISAG^hi^ T cells relative to PBMCs and all T cells remained significant after addition of these independent controls, despite a single outlier control sample with very high levels of ISAG^hi^ T cells (supplementary Figure S2). We recently reported a reduction in peripheral non-classical monocytes in familial tauopathy [5]. Comparing the familial tauopathy and EOAD datasets, we found that non-classical monocytes are not reduced in EOAD, and ISAG^hi^ T cells are not expanded in familial tauopathy (supplementary Figure S3), suggesting distinct peripheral immune responses in sporadic EOAD and primary familial tauopathy.

### 3.4 Expansion of proliferating natural killer cells in EOAD

Previous single-cell analyses have revealed additional PBMC types temporally associated with heightened type I IFN signaling. In particular, a rare NK cell subpopulation that expresses markers of proliferation has been shown to significantly expand after vaccination with an experimental HIV vaccine [8]. This expansion coincides with heightened type I IFN signaling in myeloid cells [8], which we also observe in EOAD (see below). After mapping the EOAD PBMC dataset onto a large, well-characterized multimodal PBMC CITE-seq dataset [8], we identified the proliferating NK cell cluster and, remarkably, observed significant expansion of this rare subpopulation in EOAD, specifically in female cases (supplementary Figure S4A). In addition, differential expression analysis confirmed significant enrichment for gene ontology (GO) and reactome terms related to IFN signaling and antiviral response within the primary NK cell cluster in EOAD (supplementary Figure S4B). These findings provide additional corroborative evidence, via a population of innate lymphoid cells, that EOAD is characterized by heightened peripheral IFN signaling.

### 3.5 Differential expression analysis of PBMC subsets in EOAD

Differential expression analysis revealed a high number of differentially expressed genes (DEGs) in classical and non-classical monocytes in EOAD, relative to controls (supplementary Figure S5A, supplementary Table S2). Remarkably, we found that, on average, ~19% of the significantly upregulated genes across all clusters (excluding those with fewer than 10 upregulated DEGs) were also ISAG^hi^ T-cell marker genes (i.e., IFN response genes; supplementary Figure S5A). GO analysis of the genes upregulated in CD4 T-cell clusters and myeloid cell clusters revealed significant enrichment for terms such as ‘IFN α/β signaling’ and ‘response to virus’ (supplementary Figure S5B and C). In EOAD, we therefore observe both expansion of a CD4 T-cell subpopulation expressing very high levels of genes associated with IFN signaling and concomitant upregulation of many of the same genes across additional lymphoid and myeloid cell types.

### 3.6 Validation of upregulated ISAG^hi^ T-cell marker genes via ddPCR

To validate our primary scRNA-seq findings, we magnetically isolated CD4 T cells from an additional cohort of EOAD cases and control participants. A droplet digital (dd)PCR-based validation assay indicated highly efficient isolation of CD4 T cells (supplementary Figure S6). Reasoning that increased expression of specific ISAG^hi^ T-cell marker genes from isolated CD4 T cells would be consistent with expansion of ISAG^hi^ T cells as well as ISAG upregulation, we performed ddPCR for ISAG^hi^ marker genes *MX1* and *IFI6* (Figure 2A). Cases and controls in the ddPCR cohort had similar average ages (Table 1; see also supplementary Methods document), excluding age as an explanatory factor. Strikingly, ddPCR confirmed increased *MX1* and *IFI6* expression in CD4 T cells from EOAD cases (Figure 2B). Increased *MX1* was observed across two independent ddPCR batches and was driven by females (Figure 2C).

**Figure 2.**
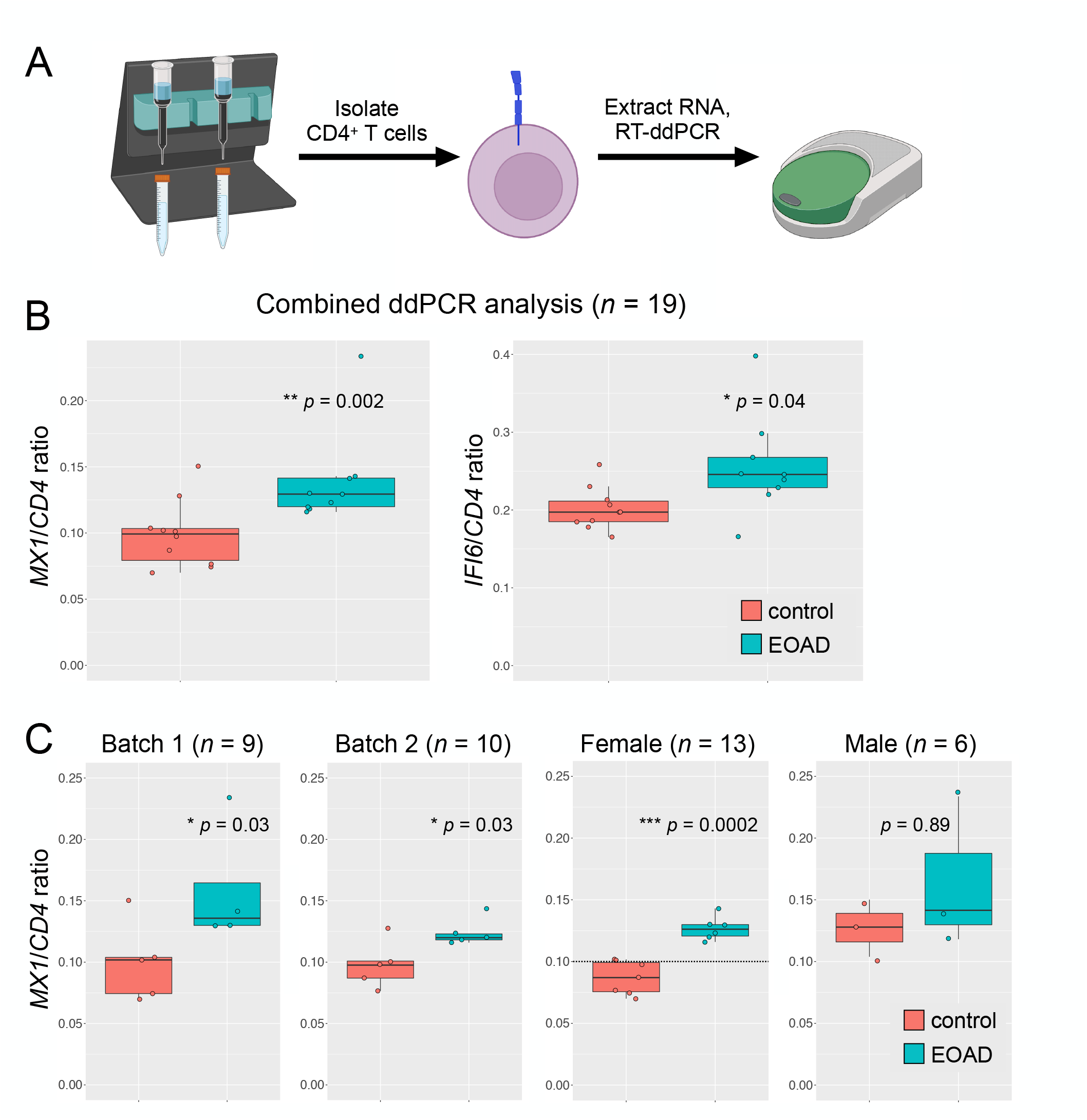
ISAG^hi^ T-cell marker gene expression is increased in CD4 T cells in EOAD. **A**, CD4 T cells were magnetically isolated from PBMCs and RNA was extracted; gene expression was determined by RT-ddPCR. **B**, Expression of *MX1* and *IFI6* was significantly increased in CD4 T cells from EOAD cases relative to cognitively normal controls (*P* = 0.002 and *P* = 0.04, respectively). **C**, *MX1* expression was significantly increased in two independent RT-ddPCR batches (*P* = 0.03, both batches). The increase in *MX1* expression observed in EOAD was driven by females (*P* = 0.0002). *CD4* was used as a reference gene.

### 3.7 Secondary analysis of CSF leukocytes in late-onset MCI/AD

Secondary analysis of a well-known CSF leukocyte dataset [3] revealed that ISAG^hi^-like T cells, although detected, were not expanded in the CSF in late-onset MCI/AD (Figure 3A through C). Strikingly, however, differential expression analysis revealed robust upregulation of *MX1* within CSF ISAG^hi^-like T cells in MCI/AD (Figure 3D). Moreover, functional enrichment analysis of the genes upregulated in MCI/AD (relative to healthy controls) across all CSF CD4 T cells revealed highly significant enrichment for terms such as ‘IFN α/β signaling’ and ‘response to virus’ (Figure 3E). In addition, analysis of upregulated DEGs from individual CSF clusters revealed similar enrichment for IFN signaling terms across multiple CD4 T-cell clusters as well as two innate immune clusters (Figure 3F). Collectively, these results suggest that, although *expansion* of ISAG^hi^ T cells may be specific to EOAD, upregulation of the same IFN signaling pathways in CSF CD4 T cells is conserved in late-onset MCI/AD.

**Figure 3.**
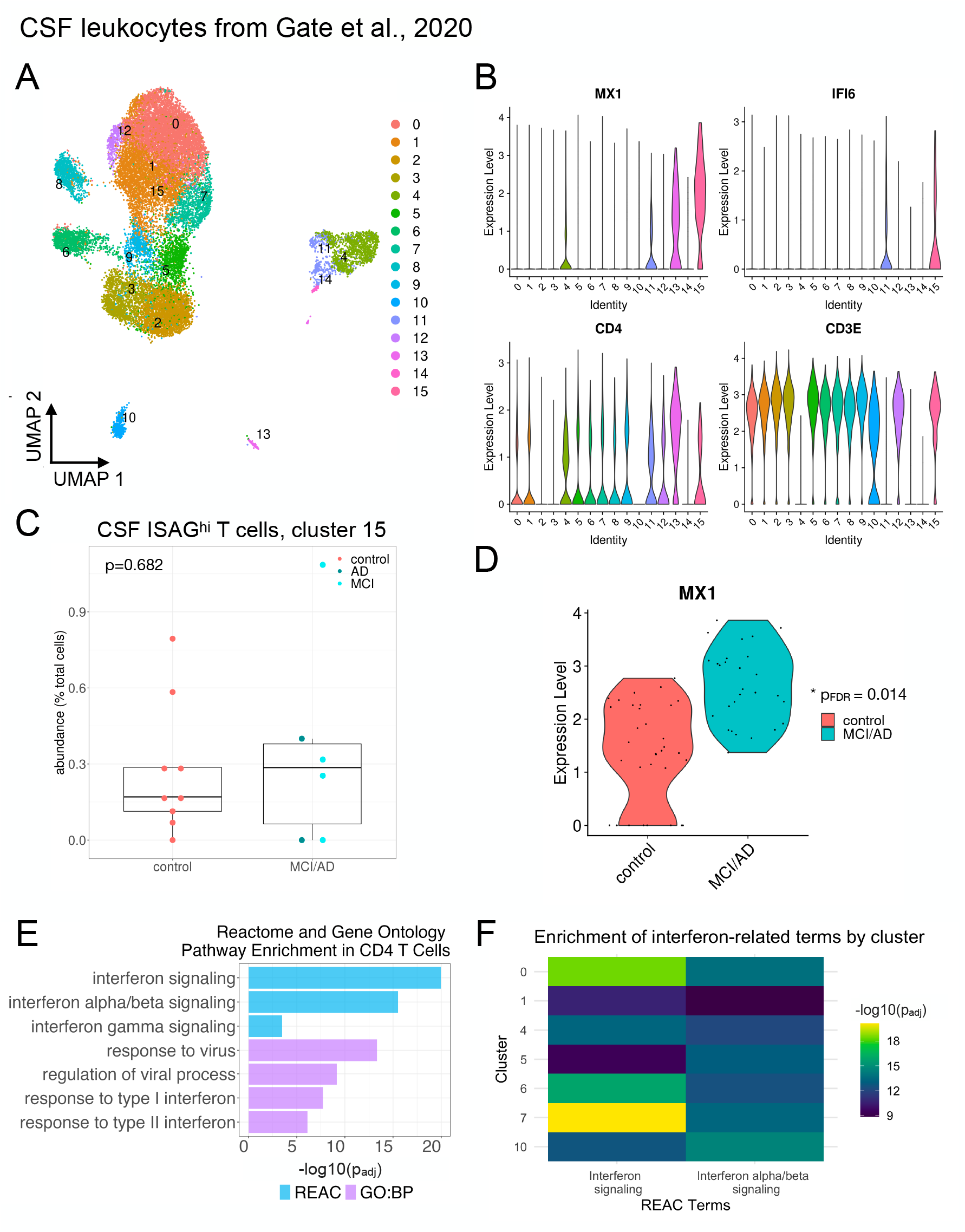
Heightened IFN response signatures in CSF T cells in MCI and AD. Publicly available data from Gate et al., 2020 [3] were downloaded from GEO (GSE134577) and analyzed as described. A UMAP plot (**A**) shows the distribution of CSF immune cells. **B**, Violin plots show that CSF cluster 15 harbors ISAG^hi^-like cells expressing high levels of *MX1* and *IFI6* along with *CD4* and *CD3E*. **C**, Quantification of CSF cluster 15 reveals lack of expansion in late-onset MCI/AD (*P* = 0.682). **D**, *MX1* expression is significantly increased (*P*_*FDR*_ = 0.014) in CSF ISAG^hi^-like T cells in MCI/AD relative to healthy controls. **E**, Functional enrichment analysis of the genes upregulated (*P*_*FDR*_ < 0.05 and log_2_ fold-change > 0.1) across all CSF CD4 T-cell clusters (0, 1, 5, 6, 7, 8, 9, 12, and 15) reveals significant enrichment of IFN and antiviral response pathways in MCI/AD. GO biological process (BP) and reactome databases were used. **F**, Analysis of significantly upregulated genes from individual CSF immune cell clusters revealed significant enrichment for IFN signaling in individual CSF CD4 T-cell clusters (0, 1, 5, 6, and 7) as well as monocyte and NK cell clusters (4 and 10, respectively).

### 3.8 Dysregulation of IFN signaling genes in a mouse model of AD

To assess the relevance of heightened type I IFN signaling in the brain, we asked whether ISAG^hi^ T-cell marker genes are dysregulated in the hippocampus and cortex in the *APP*swe x *PSEN1*.M146V (TASTPM) mouse model [17] of familial EOAD. In the TASTPM model, we observed marked upregulation of many ISAG^hi^ marker genes, particularly in the hippocampus (supplementary Figure S7A and B). These results confirm the relevance of dysregulated type I IFN signaling in the brain in a model of EOAD-like amyloidosis.

## 4 DISCUSSION

In this study, we found evidence for a unique peripheral immune signature in EOAD. Our findings complement and expand upon existing evidence of diverse T-cell signatures in other forms of AD [3,18], additional neurodegenerative diseases [4], and during aging [6]. Our study is limited by the relatively small sample sizes that are characteristic of scRNA-seq experiments. Future studies in diverse EOAD cohorts from additional recruitment sites will be needed to confirm the broad relevance of our findings to EOAD. In light of recent findings suggesting that (i) herpes zoster vaccination may be causally associated with reduced dementia risk in women [19]; (ii) viral encephalitis exposure markedly increases risk for AD [20]; and (iii) the ISAG^hi^ T cells increased in EOAD bear striking resemblance to antiviral CD4 T cells expanded in CSF in viral encephalitis [7], our findings raise the intriguing possibility that AD is characterized by an IFN-driven, antiviral-like T-cell response in both peripheral blood and CSF.

Recent work in chronic graft-versus-host disease (cGVHD) has revealed a population of naive CD4 T cells expressing an IFN response signature including *IFI6, ISG15, IFI44L, STAT1*, and *MX1* [21], genes which are all top markers of the ISAG^hi^ T cells described here and elsewhere [15], and of antiviral CD4 T cells expanded in viral encephalitis [7]. This population is strikingly enriched on day 100 post-transplant in patients who ultimately developed cGVHD, relative to those who remained tolerant. This finding strongly suggests that ISAG^hi^-like CD4 T cells contribute to tissue destruction in the context of cGVHD and may therefore also possess pathogenicity in other contexts, including AD. Future functional studies will be required to directly assess the pathogenic potential of ISAG^hi^ T cells in neurodegenerative disease models.

Complementing these insights, transcriptomic analyses have demonstrated heightened expression of genes linked to IFN signaling associated with the development of post-traumatic stress disorder (PTSD) [22,23]. Intriguingly, upon examination of detailed clinical notes for our additional control scRNA-seq samples, we found that the outlier with markedly higher levels of ISAG^hi^ T cells (supplementary Figure S2) had been previously diagnosed with PTSD. These findings suggest that the relevance of ISAG^hi^ T cells may extend beyond neurodegenerative disease to neuropsychiatric conditions as well.

In addition to our findings involving T cells, we also found evidence for heightened IFN signaling in myeloid cells and NK cells. Indeed, augmented type I IFN signaling in myeloid cells has previously been associated with expansion of proliferating NK cells [8]. Although these observations were originally reported in the context of vaccination, we observed both of these phenomena in EOAD. Given that type I IFN signaling has been shown to promote NK cell expansion and survival [24], our findings here not only support these prior observations but also provide additional evidence for heightened peripheral IFN signaling in EOAD in PBMC types beyond T cells.

Our findings build upon prior research that has found increasing evidence for heightened T-cell infiltration into the brain in AD [3,25] and AD models [25,26]. In addition, recent work in AD, primary tauopathy, and related model systems has implicated dysregulated IFN signaling pathways in microglia [25,27–32] and brain barrier tissue [33], indicating that IFN signaling is implicated not only in peripheral and CSF immune cells–as shown here–but also at the blood– CSF barrier and in brain-resident myeloid cells. Indeed, the heightened expression of ISAG^hi^ marker genes in the brain in the TASTPM model that we confirmed here may be mediated primarily by changes in microglial gene expression. Collectively, our novel findings, coupled with this prior body of work, suggest the importance of heightened IFN signaling in PBMCs, CSF immune cells, brain border tissue, and brain parenchyma, which may be mediated by distinct cellular populations in each compartment. Future work should focus on identifying the functional and compartment-specific roles of these IFN-responsive cells in neurodegenerative disease.

## Acknowledgments

The cartoon in Figure 2 was created using BioRender.com.

## Consent Statement

All participants or their surrogates provided written informed consent prior to study participation; all aspects of the studies described here were approved by the UCSF institutional review board.

