## Supplementary methods and figure legends for "Expansion of highly interferon-responsive T cells in early-onset Alzheimer’s disease"

### SUPPLEMENTARY ONLINE CONTENT

### SUPPLEMENTARY METHODS

#### Study Participants

Study participants were recruited as part of ongoing studies of Alzheimer's disease (AD) and related neurodegenerative diseases at the University of California, San Francisco (UCSF) Memory and Aging Center (MAC). All participants or their surrogates provided written informed consent prior to study participation, and all aspects of the studies described here were approved by the UCSF institutional review board (IRB). Sixteen individuals ($n$ = 8 early-onset Alzheimer's disease [EOAD] cases and $n$ = 8 cognitively normal controls) participated in the single-cell RNA sequencing (scRNA-seq) study. Nineteen individuals ($n$ = 9 EOAD cases and $n$ = 10 cognitively normal controls) participated in the droplet digital (dd)PCR study. To ensure replicability across methodologies, a subset of 4 EOAD cases was included in both studies. EOAD cases and cognitively normal controls were sex-matched, and equal numbers of female and male participants were included in the scRNA-seq study. The EOAD group was significantly older than the control group in the scRNA-seq study ($p$ = 0.0004), while ages did not significantly differ between cases and controls in the ddPCR study ($p$ = 0.37), as assessed using unpaired t-tests. Demographic information for the scRNA-seq and ddPCR study participants is included in Table 1.

#### Cell Isolation

Human peripheral blood mononuclear cells (PBMCs) were obtained from study participants at the UCSF MAC. Blood samples were collected in yellow-cap ACD Vacutainer tubes (BD Biosciences) and processed within five hours of collection, as described previously [1]. PBMCs were isolated by Ficoll density gradient sedimentation using Lymphosep separation medium (MP Biomedicals), washed with $Ca^{2+}$- and $Mg^{2+}$-free PBS (ThermoFisher), and treated with red blood cell lysis buffer (Biolegend). PBMCs were then washed again with PBS and diluted to a density of 1.5 x $10^6$ cells/ml in freezing medium composed of 10% DMSO in FBS and immediately frozen at -80°C. Samples were transferred to liquid nitrogen for long-term storage after two weeks of storage at -80°C. All PBMC samples used in our primary analyses were cryopreserved.

### Single-Cell RNA-seq

PBMCs were thawed and prepared for scRNA-seq using the Chromium Single Cell 3' v3 kit according to the manufacturer's instructions (10x Genomics). Samples were processed in two separate batches of eight samples each, with four EOAD cases and four controls included in each batch. To minimize the potential for batch effects, each batch contained equal numbers of samples from female and male participants. After sample thawing, counting, and dilution, PBMCs underwent standard 10x processing, 3' gene expression library construction steps, and next-generation sequencing at the UCSF Genomics CoLab and Institute for Human Genetics.

### Sequencing Data Processing

For each of the two batches, single-cell 3' libraries generated from eight samples were pooled and sequenced on one lane of a NovaSeq S4 flow cell. Raw sequencing reads were aligned to GRCh38-2020-A, and feature-barcode matrices were generated using Cell Ranger version 7.1.0 with intronic reads excluded.

### Quality Control

We obtained a total of $7.4 \times 10^9$ reads and detected ~260,000 cells across the two independent 10x and sequencing batches, yielding a moderate sequencing depth [2,3] of ~30,000 mean reads/cell. We detected ~5,300 median UMI counts/cell and ~1,600 median genes/cell (supplementary Table S3). There were no significant differences in the number of cells captured per sample, the number of reads per sample, or the mean read depth per sample when comparing the EOAD group to the control group. Subsequent quality-control (QC) and downstream analysis steps were performed using Seurat v4.3.0.1 [4,5]. QC filtering was applied to individual-sample feature-barcode matrices and consisted of the following steps: (i) genes detected in < 10 cells were removed; (ii) cells with ≤ 500 detected genes were removed; (iii) cells with ≤ 500 counts and those with ≥ 20,000 counts were removed; (iv) cells with mitochondrial mapping percentages ≥ 10 were removed; (v) doublets were identified and removed using DoubletFinder v2.0.3 [6,7] using the recommended parameter settings. After stringent QC filtering, ~182,000 cells remained for downstream analysis (supplementary Table S4).

### Clustering

After QC, we performed the following additional processing steps: (i) we applied sctransform v2 regularization [8,9] at the individual-sample level, including mitochondrial mapping percentage as a covariate [8,10], to minimize variability due to differences in sequencing depth between samples; (ii) the 16 individual samples were integrated with FindIntegrationAnchors and IntegrateData, specifying ‘sctransform’ as the normalization method and canonical correlation analysis (CCA) as the reduction. Subsequently, PCA was performed followed by uniform manifold approximation and projection (UMAP) reduction using the first 30 PCs; clustering was performed using a resolution parameter of 0.5. This resulted in the generation of 19 clusters that were annotated via multimodal reference mapping to a large, well-characterized human PBMC dataset [5]. Sub-clustering was performed by subsetting all T cells (as identified by multimodal reference mapping) followed by PCA, UMAP reduction using the first 30 PCs, and clustering using a resolution of 0.5. To integrate additional cognitively normal control samples from a previously published study [1] into the dataset, we performed the same analyses outlined above using a resolution of 0.4.

### Cluster Proportionality

Cluster proportions (expressed as % of total PBMCs) were determined for individual samples by dividing the number of cells in a given cluster by the total number of cells in all clusters (after QC filtering) for each individual. To determine cluster abundance relative to all T cells or all CD4 T cells, a similar calculation was performed, using as the denominator all cells annotated as T cells or CD4 T cells by multimodal reference mapping. Differences in cluster proportionality were initially assessed visually for all clusters according to disease status. Only the interferon (IFN) signaling-associated gene (ISAG)$^{hi}$ T-cell cluster (cluster 15) showed a clear difference in proportionality between cases and controls. Statistical significance for cluster 15 proportionality, as well as ISAG$^{hi}$ T-cell subcluster 11 proportionality, was determined by linear modeling, covarying for age and sex. To test for significant differences in ISAG$^{hi}$ T-cell relative abundance in the dataset containing seven additional cognitively normal controls, the cluster proportions were $\log_2$-transformed prior to analysis to minimize the effect of the single outlier control sample.

Differential Expression Analysis

Differential expression (DE) analysis was performed using limma [11–14] on individual clusters and sctransform v2-normalized data, with disease status as the contrast and age, sex and batch as covariates (as in [1]). The publicly available CSF dataset (GSE134577) was analyzed essentially in the same manner, covarying for age and sex. To account for multiple testing, a false discovery rate-corrected *p*-value ($p_{FDR}$) < 0.05 was considered statistically significant, and only genes with estimated absolute $\log_2$ fold-changes (LFC) > 0.1 were considered differentially expressed.

CD4 T cell isolation

CD4 T cells were isolated from PBMCs using the magnetic-activated cell sorting (MACS) human CD4 T cell isolation kit (Miltenyi). The purity of the isolated CD4 T cells was assessed using a droplet digital (dd)PCR assay that measured the enrichment of the *CD4*/*CD8A* transcript ratio relative to the starting PBMC population. This assay indicated an ~4,800-fold enrichment of the *CD4*/*CD8A* ratio in the isolated CD4 T cells relative to starting PBMCs, indicating successful isolation of CD4 T cells.

RNA Extraction

RNA was extracted from CD4 T cells using the RNeasy Micro Kit (Qiagen) and isolated RNA was quantified and its quality was assessed using the RNA 6000 Pico Bioanalyzer kit (Agilent). CD4 T-cell RNA samples had RNA integrity number (RIN) values ranging from 8.5–10, indicating high-quality RNA (supplementary Table S5) [15].

Droplet Digital (dd) PCR Cohort

For ddPCR experiments, we isolated CD4 T cell RNA from 19 participants. In the first ddPCR batch, we selected 5 independent controls and 4 EOAD cases with a clear expansion of ISAG$^{hi}$ T cells, as assessed by scRNA-seq. These cases were selected to determine the suitability of ddPCR for assessing the ISAG$^{hi}$ T-cell phenotype. The second ddPCR batch consisted of an additional 5 independent cases and 5 independent controls.

ddPCR Procedure and Analysis

For primary analyses, two ng of total RNA was used for single-tube reverse transcription (RT) and ddPCR using the One-Step RT-ddPCR Advanced kit (Bio-Rad). For validation of the CD4 T-cell isolation procedure, 30 ng of RNA was used to ensure that the low *CD8A* signal could be measured over background in the isolated CD4 T cells. Droplets were generated and subsequently analyzed using the QX100 system (Bio-Rad) at the UCSF Center for Advanced Technology (CAT). Reactions were prepared and run essentially according to the manufacturer's instructions. For steps in which a temperature range was specified, we used the following parameters: RT was performed at 50°C, annealing/extension occurred at 55°C, and samples were held at 12°C in the C1000 thermocycler (Bio-Rad) prior to analysis on the droplet reader. To confirm specificity, we ran the following control reactions: wells lacking RNA but containing all other components and wells lacking reverse transcriptase but containing all other components. PrimePCR ddPCR Gene Expression primer-probe mixes coupled to FAM or HEX (Bio-Rad) were used to amplify specific genes. For the analysis of ddPCR data, we performed linear modeling to assess whether disease status predicted differences in gene expression while covarying for age and sex. $Log_2$-transformed absolute concentration data for *MX1* and *IFI6* (normalized to *CD4* as the reference gene) were used for analyses that assume normality, while non-transformed data are displayed in the plots. Differences were considered significant at $p \leq 0.05$.

Secondary Analysis of Publicly Available CSF Leukocyte Dataset

Publicly available human CSF leukocyte scRNA-seq data were downloaded from GEO (accession GSE134577) [16]. The original cohort consists of samples from 18 individuals ($n$ = 9 healthy older controls, $n$ = 5 participants with mild cognitive impairment (MCI), and $n$ = 4 participants with AD). The mean (SD) ages of each group are as follows: controls, 73.1 (6.5); MCI, 69.4 (13.6); AD, 70.5 (11.5). For additional information, see Gate et al., 2020 [16]. The downloaded counts data was processed and analyzed as described above for our PBMC scRNA-seq data. We used the marker genes *MX1*, *IFI6*, *CD3E*, and *CD4* to identify a CSF leukocyte cluster corresponding to $ISAG^{hi}$ T cells. For our analyses, in order to focus on the cases with late onset, we excluded three cases that had ages at collection below 65 years. DE analysis was performed as described above for the EOAD PBMC dataset, except that cases were defined as samples with diagnoses of either MCI or AD. Genes significantly upregulated in MCI/AD with $p_{FDR} < 0.05$ and LFC > 0.1 from any CSF CD4 T-cell cluster (expressing *CD3E* and

*CD4*; clusters 0, 1, 5, 6, 7, 8, 9, 12, and 15; see Figure 3) were pooled and submitted for gene ontology (GO) analysis via g:Profiler [17]. We queried the GO biological process (BP) [18] and reactome databases [19] and selected for display significantly enriched terms related to IFN and antiviral responses. For the functional enrichment analysis of individual CSF cell clusters, we submitted upregulated DEGs from each cluster that had at least 10 upregulated genes; all such clusters, except cluster 11, showed significant enrichment for the reactome terms 'interferon signaling' and 'interferon alpha/beta signaling'.

Secondary Analysis of Mouseac Dataset

To further evaluate the expression of ISAG$^{hi}$ T-cell marker genes in a mouse model of AD-like amyloidosis, we examined genes implicated by our prior analyses in an AD model compared to wild type (WT) mice. In particular, we queried the most significantly overlapping set of ISAG$^{hi}$ marker genes between our ISAG$^{hi}$ T-cell cluster and the CSF antiviral CD4 T-cell cluster from Heming et al., 2021 [20]. Data for these analyses was obtained from the Mouseac project, which has been described in detail elsewhere [21,22]. Briefly, the Mouseac project is a study of murine brain tissue at varying ages (i.e., 8, 16, 32, and 72 weeks) in models of neurodegeneration. Samples were collected from cortex, hippocampus, and cerebellum using the TASTPM model of AD (TAS10 × TPM AD mouse models; APPswe × PS1.M146V) and WT mice [23]. Of note, both heterozygous and homozygous TASTPM mice were included in our analyses. Gene expression was measured using Illumina Ref8 v2 microarray with processing completed by the Mouseac project as previously described [21]. Briefly, expression data was $\log_2$ normalized with quantile normalization. Individual probes were excluded if the detection p-value was > 0.05 in > 50% of samples. A total of 16 ISAG$^{hi}$ marker genes were available for analysis in the downloaded dataset. ANOVA was used to analyze gene expression in comparing across age groups, gene status, and brain region. Following this, comparisons by TASTPM status were repeated within the hippocampus and cortex, the results of which are shown in supplementary Figure S7A. Differences by TASTPM status were considered significant for the hippocampus and cortex at $P = 3.1 \times 10^{-3}$, accounting for 16 genes being tested.

Additional Information

Analyses were performed in R and plots were generated using Seurat and ggplot2. For the statistical analysis of overlap between marker gene lists, we used DynaVenn [24].

### Availability of Data

The scRNA-seq dataset described here has been posted to the FAIR Data Sharing Portal within the Alzheimer's Disease Workbench, which is supported by the Alzheimer's Disease Data Initiative, and will be made accessible upon publication at: https://fair.addi.ad-datainitiative.org/#/data/datasets/single_cell_rna_seq_data_derived_from_early_onset_ad_cases_and_controls

[15] Schroeder A, Mueller O, Stocker S, Salowsky R, Leiber M, Gassmann M, et al. The RIN: an

RNA integrity number for assigning integrity values to RNA measurements. BMC Mol Biol 2006;7:1–14.

**Figure S1. Analysis of additional variables' relationships with ISAG$^{hi}$ T cells.**

**A**, Cluster 15 displays robust expression of both *CD4* and *CD3E*, indicating that it contains CD4 T cells. **B**, Estimated relative abundance of ISAG$^{hi}$ T cells (expressed as percentage of total PBMCs) was consistent across two scRNA-seq batches. **C**, *APOE* ε4 carrier status was not clearly associated with ISAG$^{hi}$ T-cell abundance in EOAD cases or controls. **D**, Increased age was not associated with increased abundance of ISAG$^{hi}$ T cells in EOAD cases or controls.

**Figure S2. Reanalysis after integration of additional cognitively normal control samples.**
Seven additional cognitively normal control samples from a prior study [1] were integrated together with the 16 samples from the present study. ISAG$^{hi}$ abundance relative to all PBMCs (top), all T cells (middle), and all CD4 T cells (bottom) was quantified. The increased abundance of ISAG$^{hi}$ T cells in EOAD remained significant relative to total PBMCs ($P$ = 0.021) and total T cells ($P$ = 0.037), while the difference relative to total CD4 T cells remained significant only after removal of the control outlier sample ($P$ = 0.014).

**Figure S3. Analysis of changes in cell-type abundance in familial tauopathy and EOAD.**

**A**, Analysis of changes in ISAG$^{hi}$ T-cell abundance in the present study and an independent study focusing on familial tauopathy revealed a specific expansion in EOAD ($P$ = 0.005). **B**, Similar analysis of changes in non-classical monocyte abundance revealed that the reduction in non-classical monocytes is specific to familial tauopathy ($P$ = 0.017).

**Figure S4. Analysis of proliferating NK cells and NK cell IFN signaling in EOAD.**
**A**, Quantitative analysis revealed a significant increase in proliferating NK cell abundance specifically in female EOAD cases compared to controls ($P$ = 0.005). **B**, Significant upregulation of IFN signaling and antiviral response pathways in NK cells from EOAD patients is

demonstrated by querying the GO BP and reactome databases with significantly upregulated genes.

**Figure S5. Differential expression analysis identifies dysregulation of ISAG$^{hi}$ T-cell marker genes in additional PBMC types in EOAD.**

**A**, Significantly upregulated genes are displayed across clusters, which are organized by PBMC type. The darker portion of each bar indicates the fraction of upregulated genes that are also ISAG$^{hi}$ T-cell marker genes. **B**, **C** (left), Volcano plots for CD4 T cells and NC monocytes (clusters 0 and 12, respectively) display downregulated (blue) and upregulated (black) differentially expressed genes (DEGs); a subset of those with $P_{FDR}$ < 0.05 and absolute LFC > 0.25 are labeled on the plots. In addition, a subset of upregulated genes with $P_{FDR}$ < 0.05 and LFC > 0.1 that are also ISAG$^{hi}$ T-cell marker genes are labeled in red. A maximum of 5 genes are labeled for each category to improve readability. **B**, **C** (right), Functional enrichment analysis of the upregulated genes ($P_{FDR}$ < 0.05 and LFC > 0.1) in CD4 T cell clusters **(B)** and in monocyte/dendritic cell clusters **(C).** There was significant enrichment of IFN and antiviral response pathways in the CD4 T cell clusters (0, 1, 6, 9, and 15) and the monocyte/dendritic cell clusters (3, 12, 14, and 16) as sourced from the GO BP and reactome databases.

**Figure S6. ddPCR validation of CD4 T-cell isolation.**

ddPCR-based quantification of the *CD4*/*CD8A* ratio from starting PBMC RNA and isolated CD4 T-cell RNA revealed an ~4,800-fold increase in the *CD4*/*CD8A* ratio upon CD4 T-cell isolation.

**Figure S7. A mouse model of familial EOAD reveals dysregulation of IFN response genes in hippocampus and cortex.**

**A**, All queried genes are displayed by heatmap in descending order of significance for TASTPM status in the hippocampus. Results are also displayed for the cortex. A total of 15 of the 16 genes queried showed significant dysregulation in mouse hippocampus. Genes not reaching significance for a given region are displayed as gray within the heatmap. **B**, The top two upregulated genes in the hippocampus (*Oas2* and *Mx2*) showed increased expression over time in TASTPM mice relative to WT mice, with TASTPM homozygotes showing greater increases relative to heterozygotes.
