## Supplementary figures and images for "Expansion of highly interferon-responsive T cells in early-onset Alzheimer’s disease"

### Supplementary Figures S1-S7

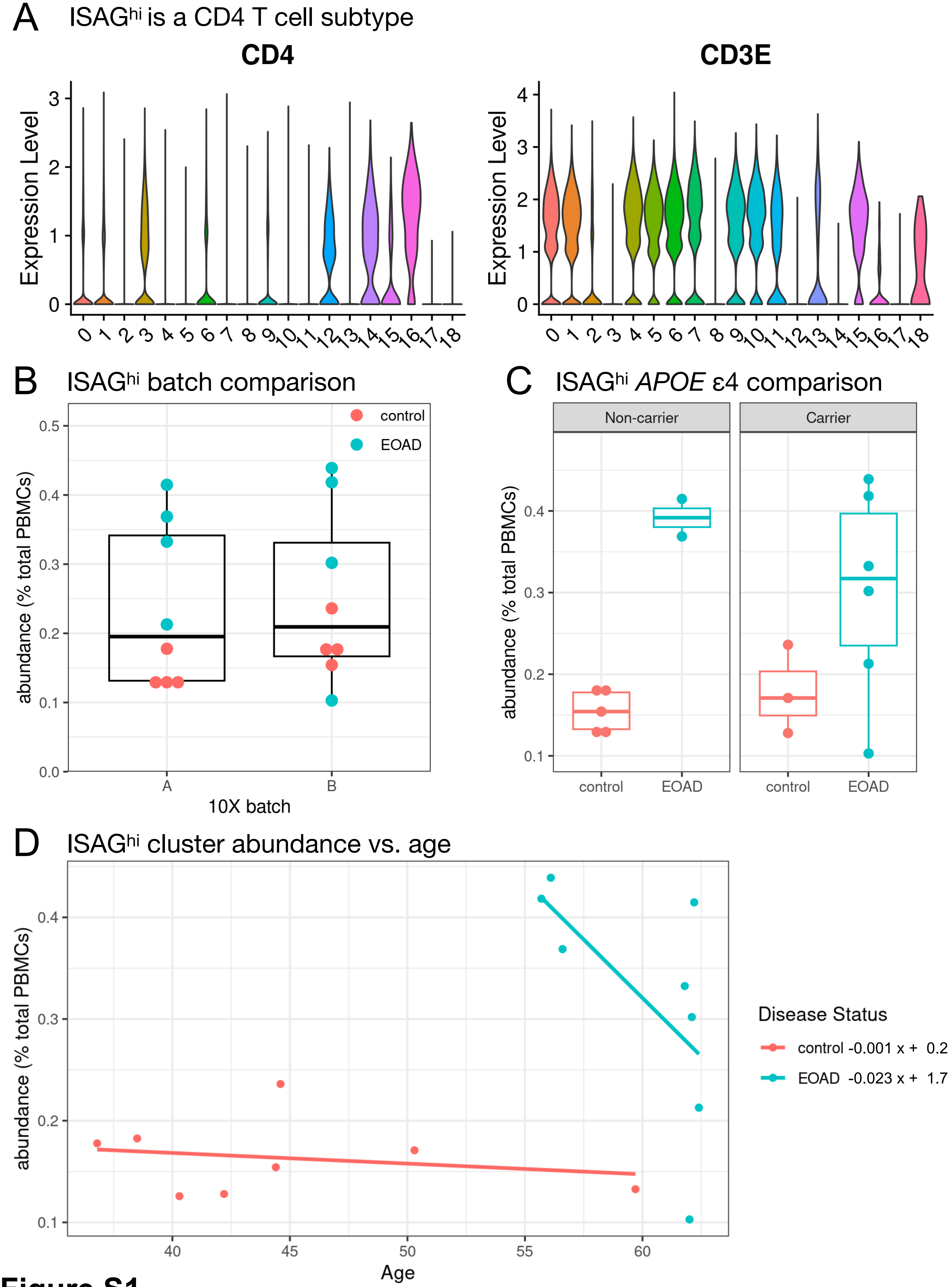

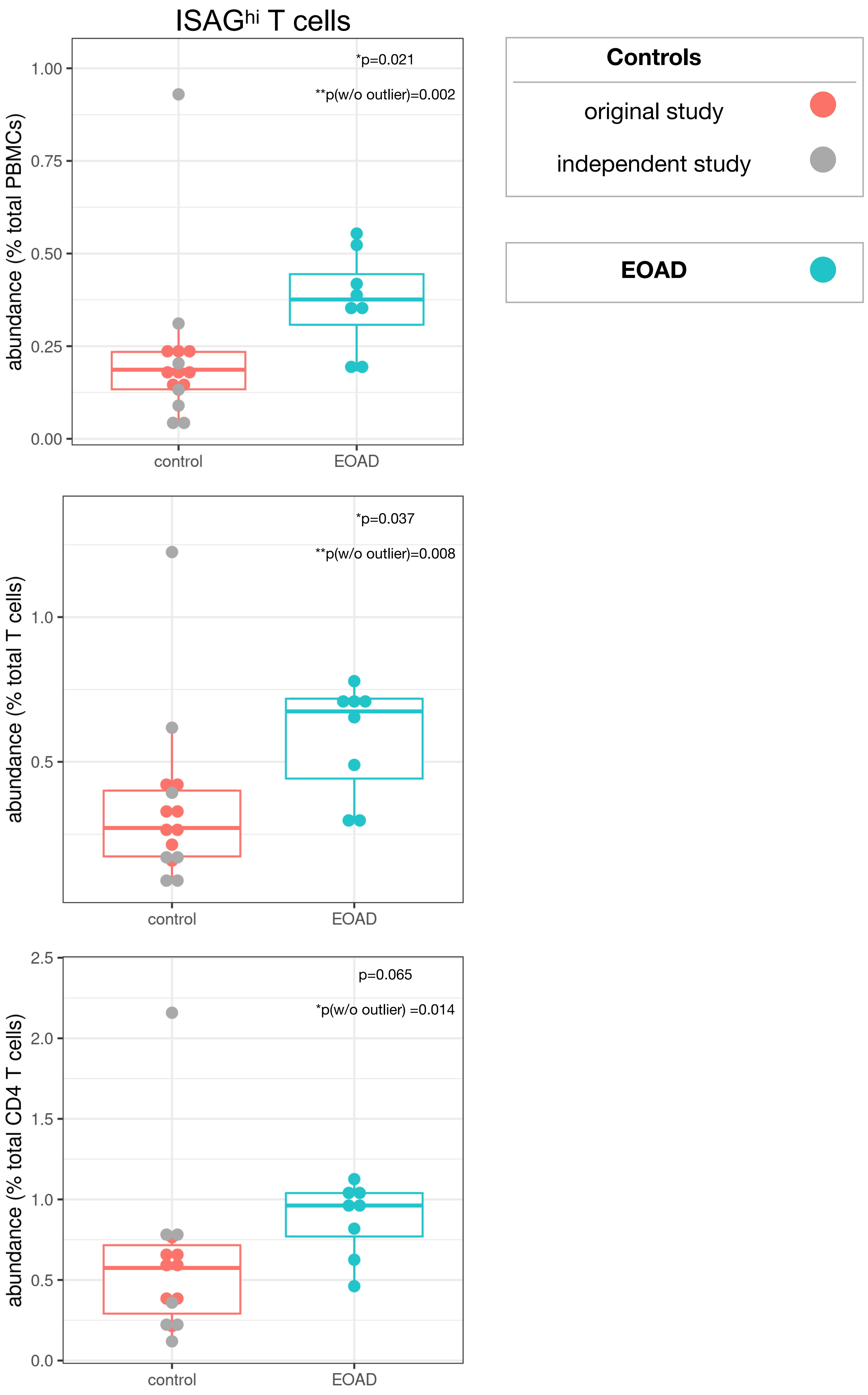

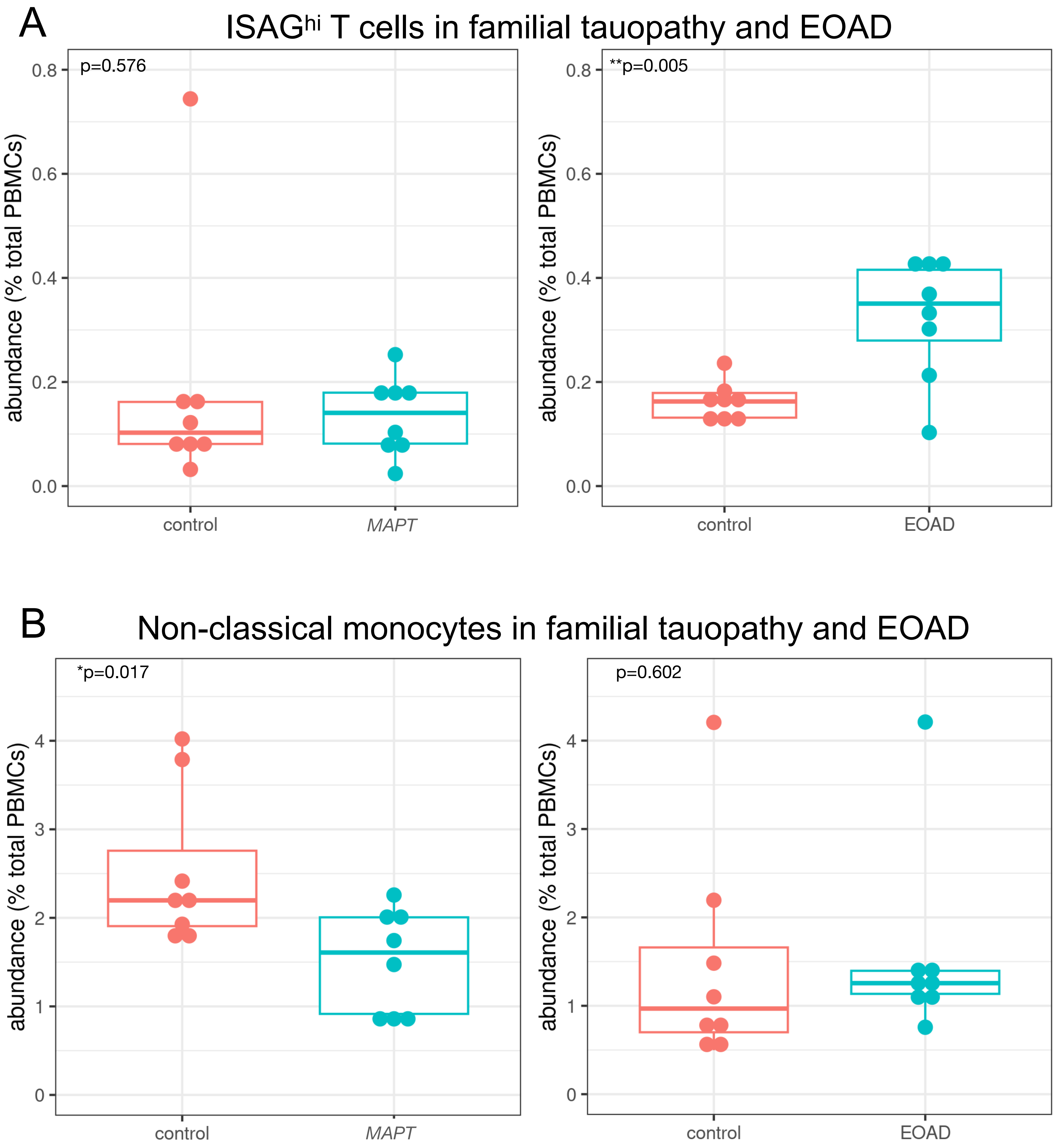

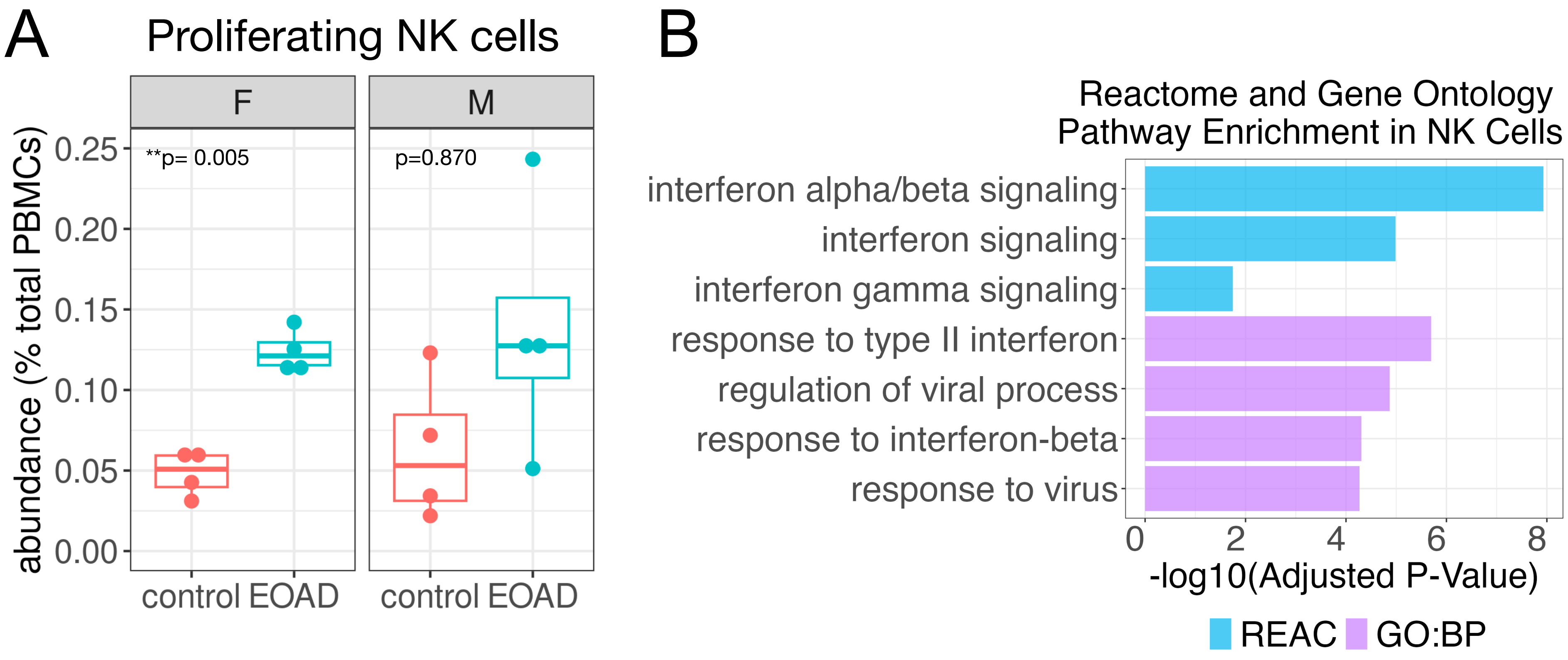

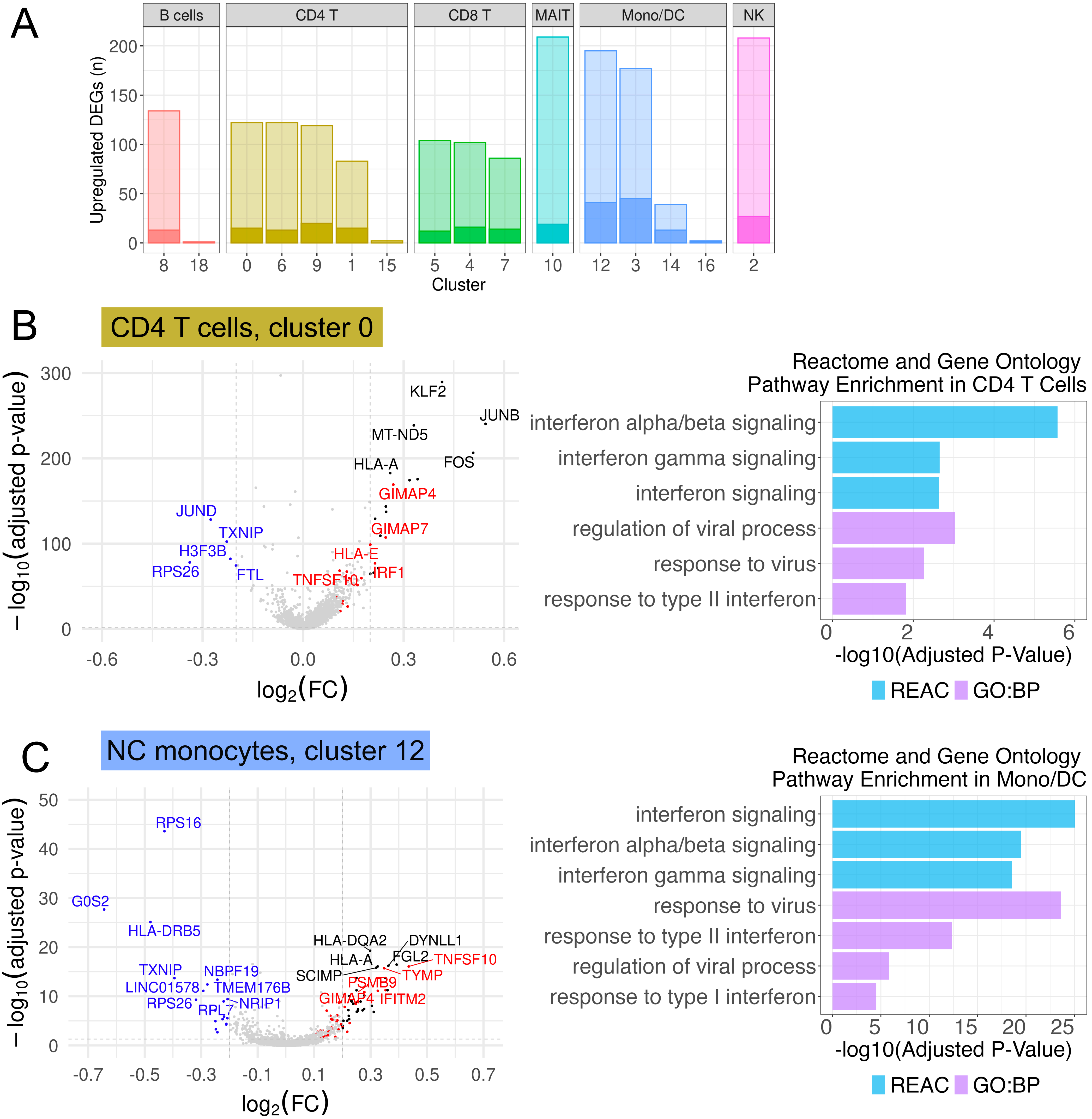

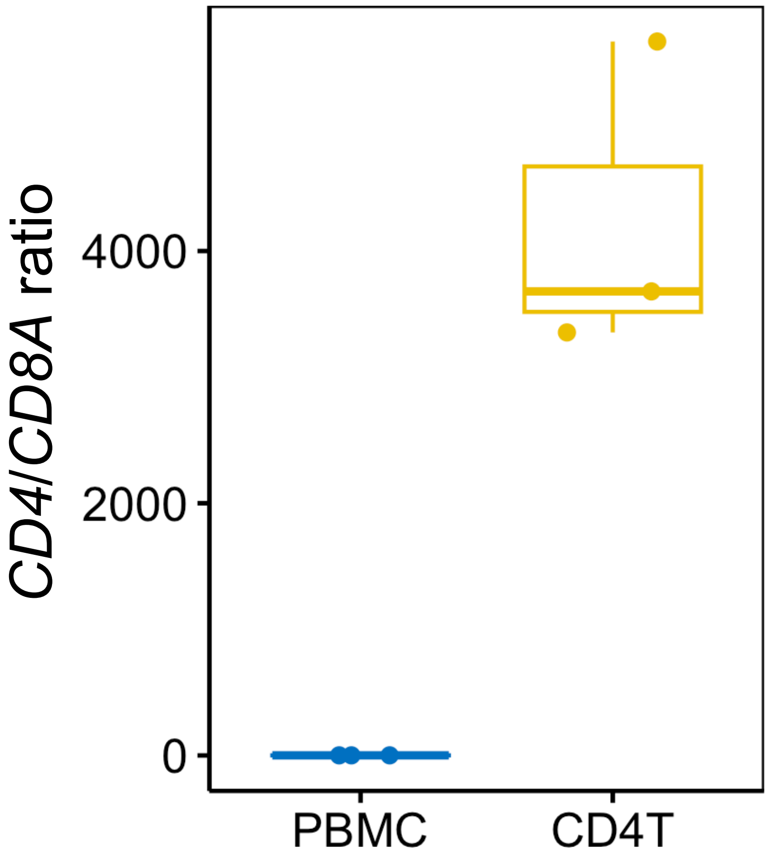


**Figure S6**

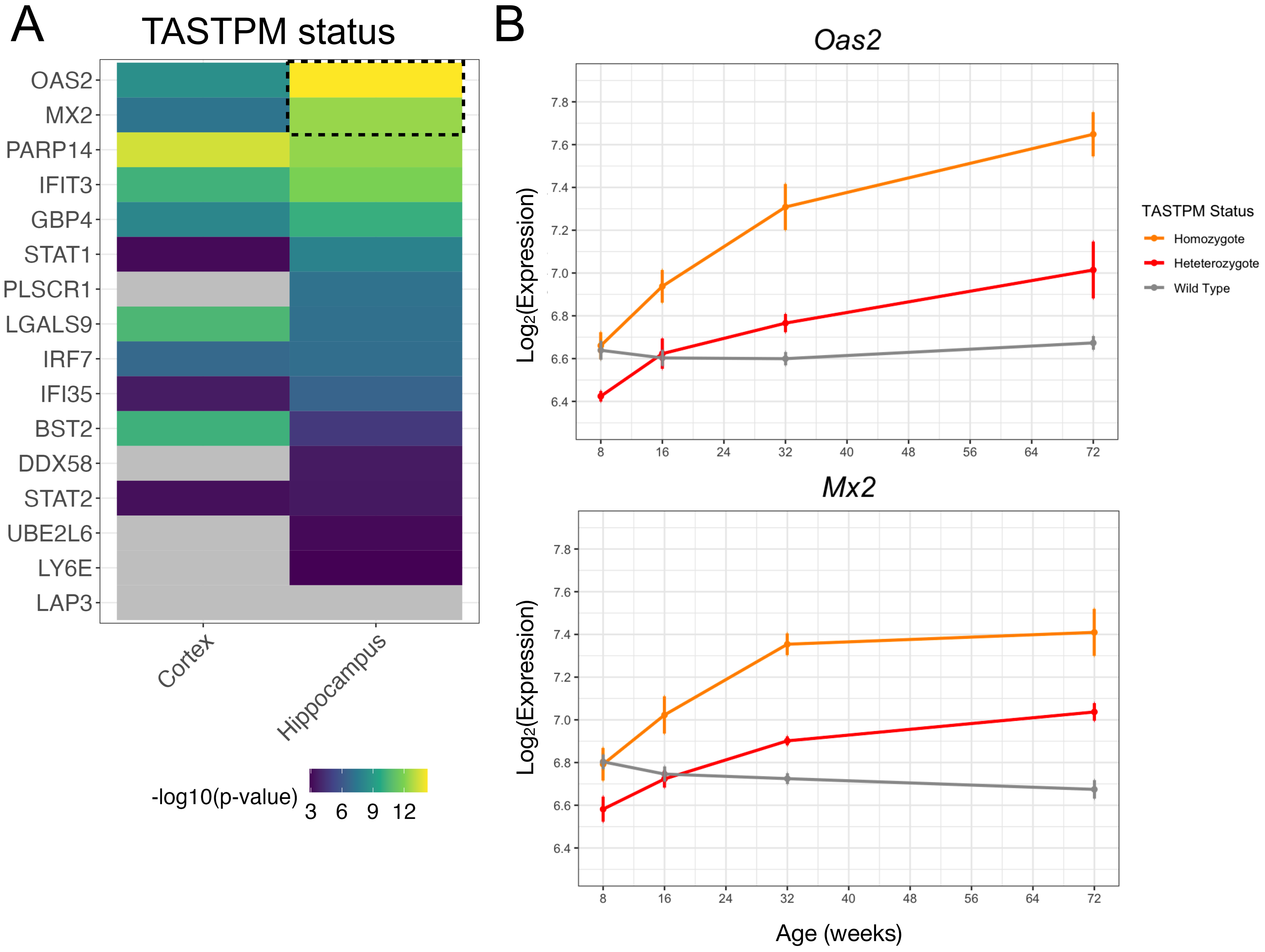
